# Practice walking on a treadmill-mounted balance beam modifies beam walking sacral movement and alters performance in other balance tasks

**DOI:** 10.1101/2022.10.30.514059

**Authors:** Evangelia-Regkina Symeonidou, Nicole M. Esposito, Roehl D. Reyes, Daniel P. Ferris

## Abstract

The goals of this study were to determine if a single 30-minute session of practice walking on a treadmill-mounted balance beam: 1) altered sacral marker movement kinematics during beam walking, and 2) affected measures of balance during treadmill walking and standing balance. Two groups of young, healthy human subjects practiced walking on a treadmill-mounted balance beam for thirty minutes. One group trained with intermittent visual occlusions and the other group trained with unperturbed vision, providing greater variation in the balance performance outcomes. We hypothesized that the subjects would show changes in sacrum movement kinematics after training and that there would be group differences due to larger improvements in beam walking performance by the visual occlusions group. We also investigated if there was any balance transfer from training on the beam to treadmill walking (margin of stability) and to standing static balance (center of pressure excursion). We found significant differences in sacral marker maximal velocity after training for both groups, but no significant differences between the two groups from training. There was limited evidence of balance transfer from beam walking practice to gait margin of stability for treadmill walking and for single-leg stance balance, but not for tandem stance balance. The number of step-offs while walking on a narrow beam had the largest change with training (partial η^2^=0.7), in accord with task specificity. Other balance metrics indicative of transfer had lower effect sizes (partial η^2^<0.5). Given the limited transfer across balance training tasks, future work should examine how intermittent visual occlusions during multi-task training improve real world functional outcomes.

## Introduction

Practicing a balance task that not only improves performance of that balance task but also transfers improvement to other balance tasks would be ideal for motor skill training and rehabilitation. Recent studies have mixed findings on the transfer of balance training. Many studies that have examined standing balance training have found little to no transfer of balance improvements to other balance tasks (Giboin et al., 2018; Kümmel et al., 2016; McCrum et al., 2017, 2018). In contrast, some studies that have looked at walking balance have shown some limited transfer to other balance tasks (Donath et al., 2017; Giboin et al., 2015). The likelihood of transfer seems to be highest if balance practice involves walking balance with self-generated transfers from single support to double support and does not have external physical perturbations disrupting balance.

One potential training method for challenging dynamic balance is walking on a narrow beam. Studies on humans walking on an overground balance beam have been useful for understanding motor skill expertise (Robertson & Elliott, 1996a; Sawers & Hafner, 2018; Sawers & Ting, 2015) and studying the effects of vision, reflex modulation, and loss of balance in walking (Llewellyn et al., 1990; Robertson et al., 1994; Robertson & Elliott, 1996b; Sipp et al., 2013). Walking on a treadmill-mounted balance beam has been used to test the effects of error augmentation and physical guidance on balance motor learning (Domingo & Ferris, 2009, 2010) and can be combined with physical perturbations, visual perturbations, or immersive virtual reality environments (Peterson, Furuichi, et al., 2018; Peterson, Rios, et al., 2018; Peterson & Ferris, 2018).

Successful locomotion across a narrow beam requires humans to control movement of their center of mass (CoM) to prevent stepping off the beam. In normal gait, humans actively maintain medio-lateral balance by controlling foot placement for each step (Arvin et al., 2016; Bruijn & Van Dieën, 2018; Kuo & Bauby, 2000). Walking on a narrow beam (e.g., 2.5 cm wide) does not allow humans to adjust foot placement in the way they do when walking overground. They do have to monitor the movement of their CoM and repeatedly shift weight from the trailing limb to the leading limb, requiring active involvement from many brain areas including prefrontal, anterior cingulate, premotor, sensorimotor, and posterior parietal cortices (Peterson & Ferris, 2018; Sipp et al., 2013).

Brief intermittent visual perturbations during beam walking practice can improve subsequent beam walking performance (Peterson, Rios, et al., 2018; Symeonidou & Ferris, 2022). Peterson et al. (2018) showed that intermittent visual rotations presented through a virtual reality headset during 30 minutes of beam walking practice led to a reduction in step-offs of 42% compared to a 9% reduction in step-offs when practicing without intermittent visual rotations (Peterson, Rios, et al., 2018). More recently, Symeonidou and Ferris (2022) used liquid crystal lens glasses to train participants with brief intermittent visual occlusions during the same beam walking paradigm. The visual occlusions group showed a 78% reduction in step-offs on the same day after training and a 60% retention two weeks later. In comparison, the unperturbed vision group only showed a 20% reduction in step-offs on the same day and a 5% retention (Symeonidou & Ferris, 2022; Symeonidou & Ferris, 2019). These studies provide strong evidence that incorporating brief reoccurring visual perturbations when learning a difficult balance task can improve subsequent performance of the same balance task. The neural mechanisms responsible for the visual perturbation effect on balance improvement are not known, but it is possible that cross-modal synchronization of sensory information processing enhances training (Axmacher et al., 2006; Fell & Axmacher, 2011; Lakatos et al., 2008; Schroeder & Lakatos, 2009; Thorne et al., 2011). From previous studies on beam walking, it is not clear how the subjects altered their body movement patterns to achieve reductions in beam step-offs with practice.

In this study, we sought to identify changes in body movement patterns related to improved beam walking performance with practice, and if there was transfer of better dynamic balance from beam walking to other balance tasks. Specifically, we wanted to determine if a single 30-minute session of practice walking on a treadmill-mounted balance beam: 1) altered sacral marker movement kinematics during beam walking, and 2) affected measures of balance during treadmill walking and standing balance. We chose sacral movement as the primary focus of our kinematic analysis for beam walking performance because it is a simple method and good approximation for estimating CoM motion during walking (Gard et al., 2004; Gordon et al., 2009; Thirunarayan et al., 1996; Yang & Pai, 2014) and treadmill beam walking in particular (Domingo & Ferris, 2009, 2010). Whole body CoM estimation would be time-consuming and requires a whole-body marker set, limiting its applicability in real-life settings. We studied two groups of subjects, one with brief intermittent visual occlusions and one with unperturbed vision, walking on a treadmill-mounted balance beam for 30 minutes. We expected the group with intermittent visual occlusions to improve more in beam walking performance than the unperturbed vision group. We hypothesized that sacral movement kinematics during beam walking would be altered after practice, and that the visual occlusions group would have greater changes from pre-test to post-test than the unperturbed vision group. As there is no clear evidence from the literature whether sacral movement would be increased or decreased with improved balance walking on a beam (Chiovetto et al., 2018; Domingo & Ferris, 2009, 2010; Huber et al., 2020), we did not *a priori* predict a direction of the change from pre-test to post-test. We also compared other metrics of balance during treadmill walking and standing balance to determine if there was any evidence of transfer from the balance beam practice to other balance measures.

## Materials and Methods

### Participants

Twenty healthy, young participants (11 males, 9 females) took part in this study. They reported no neurological, orthopedic, musculoskeletal conditions or lower limb surgeries. All participants were right-foot dominant. We assessed foot-dominance by asking them which foot they would use to kick a ball. Written, informed consent was obtained from all study participants and the study was approved by the University of Florida Institutional Review Board (IRB). The study was conducted according to the Declaration of Helsinki.

### Experimental Design

#### Beam Training

Participants performed a beam walking task with or without the presence of reoccurring intermittent visual occlusions (Fig. 1). The balance beam walking task consisted of a 3 min pre-test trial, 30 minutes of training, and a 3-min minute posttest trial. One group trained with brief intermittent visual occlusions (5 females, 5 males, age = 25.9 ± 4.5) that were presented over liquid crystal lens glasses (Senaptec Strobe, Senaptec, Oregon, 148 USA) in a periodic fashion (1.5 s occlusion and 7.5 s of clear vision) and another group (5 females, 5 males, age= 25.2 ± 4.7) trained on the beam while wearing the glasses but without any visual occlusions. The treadmill-mounted beam was 2.5 cm high and 2.5 cm wide (Domingo & Ferris, 2009; Peterson, Rios, et al., 2018; Sipp et al., 2013; Symeonidou & Ferris, 2022) and participants walked on a constant speed of 0.22 m/s (Domingo & Ferris, 2009; Peterson & Ferris, 2018; Symeonidou & Ferris, 2022).

**Figure 1.**
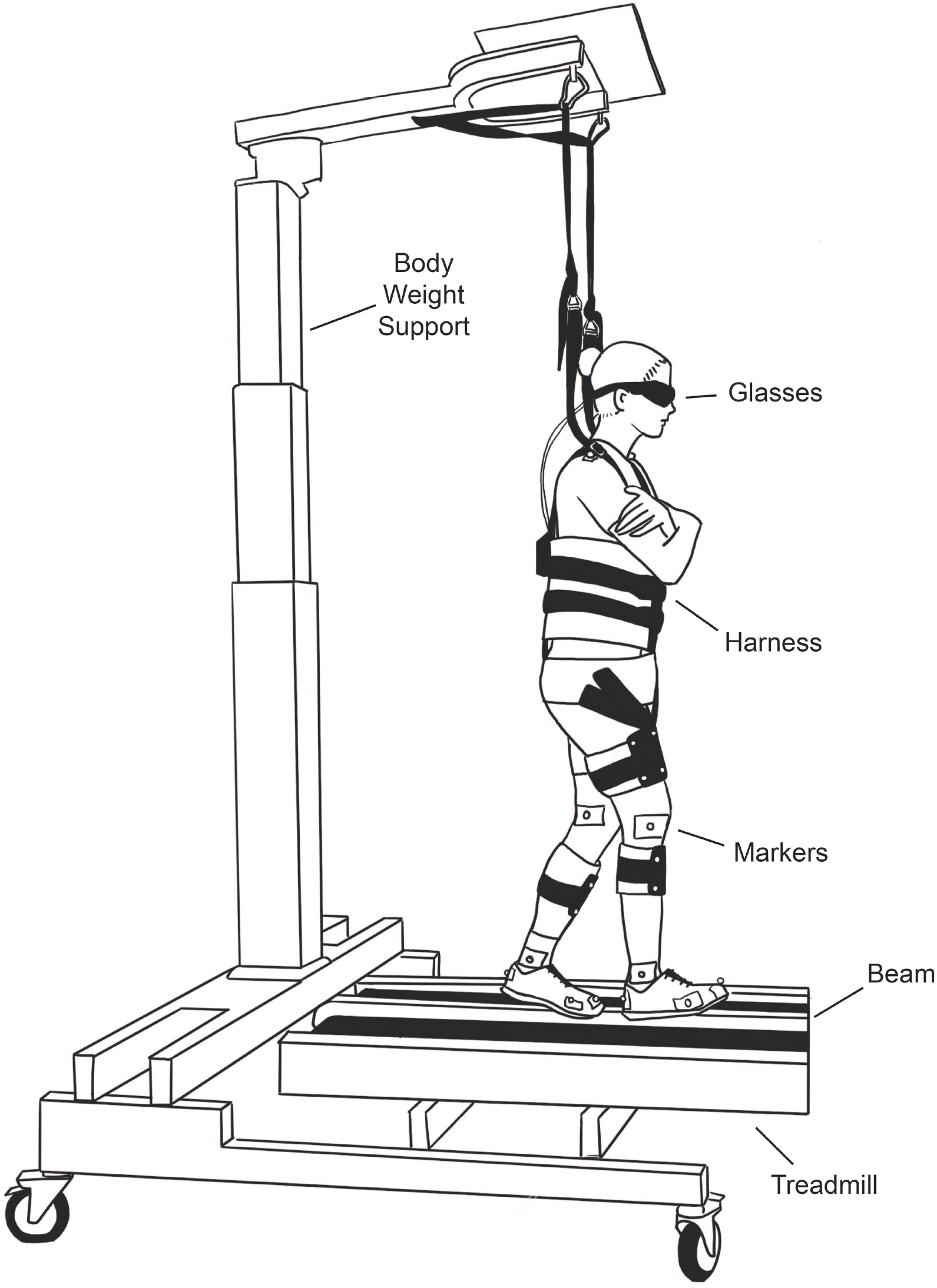
Experimental Setup. Participants performed the beam walking training on a treadmill-mounted balance beam while wearing the occlusion glasses. They were strapped for safety in a harness attached to a body-weight support system that was modified to allow for freedom of movement in the mediolateral direction. A number of reflective markers were attached to the participant’s body for motion tracking purposes.

The visual occlusions were presented in a periodic manner using a Samsung Note 10 that was running the Senpatec app (Senaptec, Oregon, USA). The occlusion cycle would start when participants stepped on the beam with both feet and ended when they stepped off. A random delay of up to 1 s was added to the start of the cycle to avoid any anticipatory effects. To start and stop the cycle we used TeamViewer and a custom MATLAB script to access the screen of the phone over a desktop monitor. Every time participants stepped off the beam, they had to count 5 steps before stepping back on (Domingo & Ferris, 2009; Peterson, Rios, et al., 2018; Peterson & Ferris, 2018). The pre-test and post-test of the balance beam walking task were performed with the glasses on but in the transparent mode for both groups. During the balance beam walking participants were able to take breaks between trials and had to take a 5-min break between the last training trial and the post-test trial.

#### Walking and Static Balance Tasks

Participants performed the walking and static balance tasks on a split-belt treadmill (Bertec, Ohio, USA) that recorded forces through embedded force plates with a sampling rate of 1000 Hz. Participants walked for 5 mins and performed a 30-sec tandem and single-leg stance before (pre), and after (post) the beam walking task (Fig. 2). The order of the tasks was counterbalanced across participants for the pre and post session. The split-belt treadmill was set at a constant speed of 0.5 m/s and participants were instructed to walk on the center of each belt in the anterior-posterior direction. For the single-leg stance task participants were instructed to stand on one belt of the treadmill and raise their left leg 10 cm off the ground. During the tandem stance task participants were instructed to stand on one belt with the right foot in front of the left foot, with the heel of the right foot touching the fingers of the left foot. For both static balance tasks, participants were asked to rest their arms at their waist, look straight, and keep this position for 30 sec.

**Figure 2.**
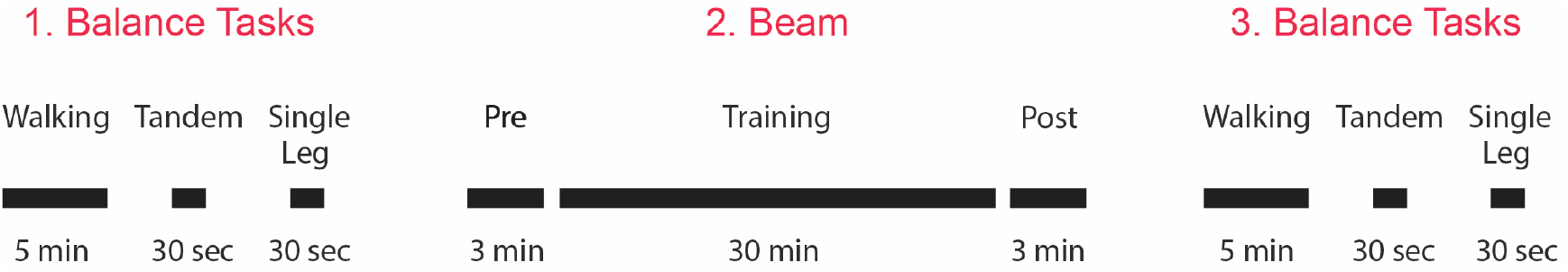
Experimental Timeline. Participants performed the standing and walking balance tasks before and after the balance beam training in randomized order: walking, tandem stance, and single-leg stance.

### Data Analysis

#### Beam Kinematics

Marker trajectories were cleaned using Motive software (Optitrack, Natural Point Inc, Corvallis, OR, USA) and processed with Visual3D (C-Motion, Germantown, MD). We low-pass filtered the marker data using a 6 Hz cut-off frequency and fourth-order Butterworth filter and determined on-beam phases using a manual threshold for the position of the toe, last metatarsal, and heel foot markers in the medio-lateral and vertical directions for each step. We used a manually adjusted, empirically determined threshold because the beam location varied slightly across subjects and sessions. A number of reflective markers (n=28) were placed on the participants’ feet, knees, hips, sacrum, back, shoulder, neck, and head, to record motion capture data (Optitrack, Natural Point Inc, Corvallis, OR) sampled at 100 Hz for all tasks (beam walking, walking, tandem stance, single-leg stance).

We calculated the sacrum kinematics when participants were on the beam in the medio-lateral, anterior-posterior, and vertical direction. We calculated the average standard deviation, as a surrogate for the CoM variability, during the pre- and post-trials for each participant and averaged the values within the two groups. To identify if the visual occlusions group employed faster corrective adjustments after the training compared to the unperturbed vision group, we looked at sacrum peak displacement and velocity for the pre- and post-trials. To do so, we calculated the highest absolute value of displacement and velocity of the sacrum marker for each trial and participant, and then averaged within groups. To investigate if participants adopted a more crouched position after training, we calculated the normalized sacrum-ankle distance for the pre- and post-trials. For each on-the-beam stance phase within each trial, we averaged the vertical component of the sacrum and ankle marker and found the distance between the two. This distance was normalized to the sacrum-ankle distance of the participant’s upright stance when off-the beam. Normalized sacrum-ankle distance values were averaged within trials and groups.

#### Walking Margin of Stability

We calculated the dynamic margin of stability to see if there was any transfer of the gained balance benefit from the balance beam training to the treadmill walking (Young et al., 2012). We estimated the CoM position using the average position of seven pelvic markers. The CoM was then used to find the extrapolated CoM (XcoM), expressed as

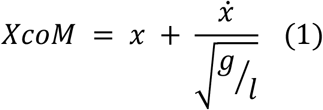

Where x is the CoM position, 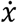 is the CoM velocity, *g* is the gravitational constant, and *l* is the equivalent pendulum length, i.e. the mean distance from the leading foot’s last metatarsal marker (left or right) to the CoM at every heel strike. The dynamic margin of stability (MOS) was then calculated as

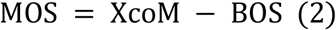

Where BOS is the boundary of the base of support. Dynamic margin of stability and base of support were both measured separately in the anterior-posterior and medio-lateral directions. The base of support was defined by the position of the leading foot marker at heel strike. Dynamic margin of stability values were calculated at heel strike for each step in the pre- and post-walking test and averaged within the whole trial. Heel strikes were detected using the force plates positioned underneath the split-belt treadmill.

#### Static Balance Measures

For the tandem stance and single-leg stance task we low-pass filtered the force plate data using a 12 Hz cut-off frequency and calculated several center of pressure (COP) measures: i) average COP sway excursion, ii) standard deviation of COP sway excursion, ii) mean COP velocity in the medio-lateral and anterior-posterior direction, and iv) area of COP sway excursion during tandem stance and single-leg stance. For all data analysis we used custom MATLAB scripts (The MathWorks Inc, Natick, MA, USA).

### Statistical Analysis

We had to exclude one participant from each group from the whole analysis due to missing force plate and marker data during the experiment (step-offs and on-beam sacral marker kinematics analysis, n=9/group). We had to exclude an additional participant from each group from the walking and static balance tasks analysis due to missing force plate and marker data during some of the relevant trials (walking and static balance kinematics analysis, n=8/group). We performed a repeated measures ANOVA with the training type (visual occlusions, unperturbed vision) as the between subjects’ factor and the test-trial (pre-test, post-test) as the within subjects’ factor for the on-beam measures i) sacrum standard deviation ii) maximum sacrum displacement ii) maximum sacrum velocity in the medio-lateral, vertical, and anterior-posterior direction, and iii) ankle-sacrum ratio. The same statistical analysis with the same between and within subject factors was performed for the beam performance in step-offs/min. Similarly, we performed a repeated measures ANOVA with the training type (visual occlusions, unperturbed vision) as the between subjects’ factor and the test-trial (pre, post) as the within subject’s factor for the i) average COP sway excursion, ii) SD of COP sway excursion, ii) mean COP velocity in the medio-lateral and anterior-posterior direction, and iv) area of COP sway excursion during tandem stance and single-leg stance. We performed the same statistical analysis with the same between and within subject factors for the i) mean margin of stability, ii) margin of stability variability in the medio-lateral and anterior-posterior direction during walking. Before deciding to use a parametric statistical test for the above measures, we checked the normal distribution of our data. Most of our data was normally distributed. Our primary and secondary outcome measure were step-offs/min and sacral marker standard deviation, which were both normally distributed. We expected limited to no changes in other balance metrics on the beam, and/or during standing and walking. Last, all significant results reported in the next section came from normally distributed data, therefore, violation of normality within some of our measures did not affect our results. We report Greenhouse-Geisser corrected values wherever the test of sphericity is violated.

## Results

Both groups experienced significantly fewer step-offs after training (F (1, 16) =31.4, p<.001, partial η^2^=.66). The visual occlusions group showed a 75% reduction in step-offs (F (1, 16) =14.9, p<.001, partial η^2^=.48, t(8)=4.9, p=.001) while the unperturbed vision group showed 23% improvement between pre-test and posttest (t(8)=2.9, p=.02). Both effect sizes (partial η^2^) of the test-trial (~0.7) and visual occlusion training (0.5) on step-offs were large (Table 1).

**Table 1.** On-Beam Performance and Kinematics. Mean and standard deviation (SD) of on-beam step-offs/min, sacrum standard deviation, maximum sacral displacement, and maximum sacral velocity in the medio-lateral (ML), vertical (Vert.), and anterior-posterior (AP) direction, as well as normalized ankle-sacrum distance before (Pre) and after (Post) the balance beam training for 8 participants/group. All kinematic measures are in cm or cm/s except for the normalized ankle-sacral distance. Significant p-values are denoted in bold with an asterisk (*).

| On-Beam Balancing |  |  |  |  |  |  |
| --- | --- | --- | --- | --- | --- | --- |
| Performance and Kinematic Measures | Unperturbed Vision | | Visual Occlusions | | p-value (partial $\eta^2$) | |
|  | Pre | Post | Pre | Post | Test-Trial | Training |
| Step-Offs/min | 15.50 | 11.97 | 9.74 | 2.43 | **0.00\*** | **0.01\*** |
| ±SD | 5.15 | 5.90 | 4.67 | 2.09 | (0.66) | (0.50) |
| Sac Standard Deviation ML | 3.79 | 3.40 | 3.32 | 2.32 | 0.09 | 0.20 |
| ±SD | 1.39 | 1.76 | 1.66 | 0.88 | (0.17) | (0.10) |
| Sac Standard Deviation Vert. | 1.28 | 1.41 | 1.17 | 1.19 | 0.42 | 0.31 |
| ±SD | 0.27 | 0.81 | 0.39 | 0.36 | (0.04) | (0.07) |
| Sac Standard Deviation AP | 6.72 | 5.51 | 5.72 | 5.10 | 0.06 | 0.18 |
| ±SD | 1.77 | 1.42 | 1.24 | 1.19 | (0.21) | (0.11) |
| Sac Max Displacement ML | 13.34 | 13.32 | 12.61 | 9.17 | 1.86 | 1.27 |
| ± SD | 2.73 | 5.39 | 4.12 | 4.03 | (0.11) | (1.40) |
| Sac Max Displacement Vert. | 4.49 | 4.84 | 4.54 | 4.46 | 0.71 | 0.80 |
| ± SD | 1.25 | 2.05 | 1.04 | 1.76 | (0.01) | (0.001) |
| Sac Max Displacement AP | 19.27 | 18.03 | 17.09 | 14.76 | 0.24 | 0.11 |
| ± SD | 5.55 | 4.91 | 4.91 | 3.42 | (0.09) | (0.16) |
| Sac Max Velocity ML | 45.24 | 45.89 | 39.76 | 34.60 | 0.40 | **0.04\*** |
| ± SD | 7.09 | 11.36 | 6.85 | 12.02 | (0.04) | (0.24) |
| Sac Max Velocity Vert. | 30.69 | 35.88 | 18.06 | 21.58 | **0.04\*** | **0.009** |
| ± SD | 7.25 | 15.61 | 6.38 | 10.38 | (0.24) | (0.35) |
| Sac Max Velocity AP | 34.51 | 37.07 | 32.49 | 33.57 | 0.06 | 4.40 |
| ± SD | 7.00 | 11.96 | 8.21 | 10.92 | (0.02) | (0.04) |
| Ankle-Sacral Distance | 100.71 | 99.00 | 100.26 | 99.59 | **0.001\*** | 0.88 |
| ± SD | 1.08 | 1.33 | 0.95 | 1.07 | (0.50) | (0.001) |

Sacral movement variability, as measured by sacral marker standard deviation, did not show any clear changes from beam walking practice. Participants of both groups showed no differences in sacral marker standard deviation between test-trials in the medio-lateral (F(1, 16) =3.3, p=.092, partial η^2^=.17), vertical (F(1, 16) =.69, p=.42, partial η^2^=.04), and anterior-posterior direction (F(1, 16) =4.2, p=.06, partial η^2^=.2) (Table 1). The training type had also no effect on sacral marker standard deviation (medio-lateral: F(1, 16) =1.8, p=.2, partial η^2^=.10; vertical: F(1, 16) =1.1, p=.31, partial η^2^=.07; vertical: F(1, 16) =.2, p=.18, partial η^2^=.11) (Table 1).

Subjects did show significant increases in maximal sacral vertical velocity after training compared to before training. There were significant main effects of test-trial (F(1,16)=4.9, p=.04, partial η^2^=.24) and training type (F(1,16)=8.7, p=.009, partial η^2^=.35) on vertical maximal sacral velocity. There was also a significant training type effect on mediolateral maximal sacral velocity (F(1,16)=5.1, p=.04, partial η^2^=.24), as the intermittent visual occlusion group had a decrease and the unperturbed vision group has a very small increase (Table 1, S1 Fig.).

Participants adopted a more crouched posture in beam walking after practice. There was a reduction in ankle-sacrum distance for both groups after training (F(1,16)=15.8, p=.001, partial η^2^=.5), but there was not a statistical difference between groups (Table 1, S2 Fig.).

For the treadmill walking data, there was only one indication of change in dynamic balance. The margin of stability in the anterior-posterior direction was larger in the post-test compared to the pre-test for both groups (F(1, 14) =5.11, p=.04, partial η^2^=.27) (Fig. 3). There was no difference between training type. There were no significant differences in margin of stability in the medio-lateral direction between test-trials or type of training. We did not detect any significant differences in margin of stability variability between test-trials or training in any direction.

**Figure 3.**
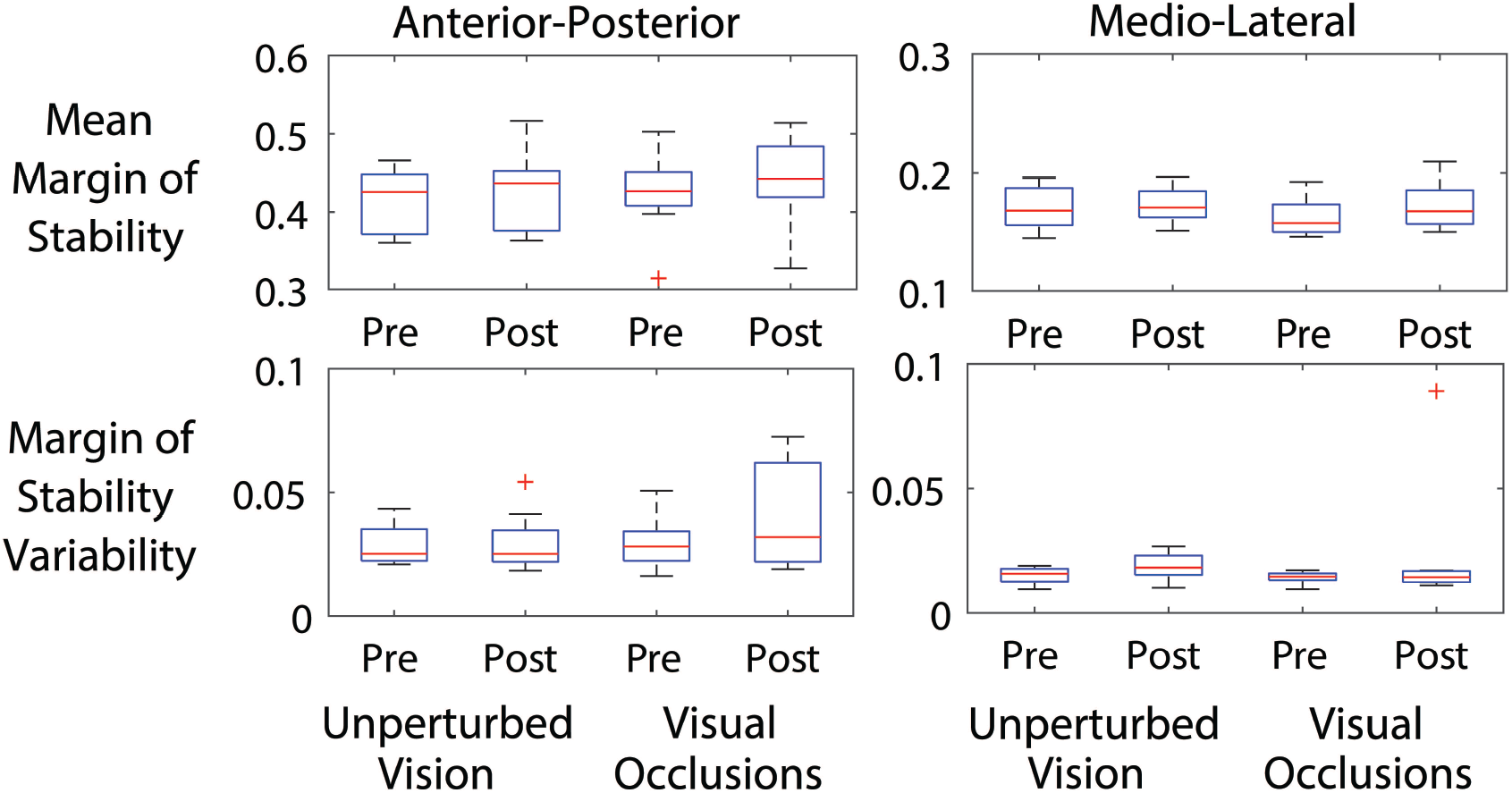
Walking Margin of Stability. Mean margin of stability and margin of stability variability for the visual occlusions and unperturbed vision group before (Pre) and after (Post) the training. There was a significant increase of margin of stability in the anterior-posterior direction between pre and post-test (F(1, 14) =5.11, p=.04, partial η^2^=.27), but no main effect of training type (visual occlusions vs unperturbed vision: F(1, 14) =.3, p=.59, partial η^2^=.021). There were no significant differences in margin of stability in the medio-lateral direction between pre and post-test (F(1, 14) =1.12, p=.31, partial η^2^=.08) or training type (F(1, 14) =.4, p=.54, partial η^2^=.03). Similarly, there were no significant differences in margin of stability variability between test-trials (anterior-posterior: F(1, 14) =1.1, p=.32, partial η^2^=.07; medio-lateral: F(1, 14) =1.1, p=.31, partial η^2^=.08) or training types (anterior-posterior: F(1, 14) =7.5, p=.4, partial η^2^=.05; medio-lateral: F(1, 14) =.4, p=.54, partial η^2^=.03) in both directions.

In the standing measures of balance, we found two indicators of training transfer. In the single-leg stance task, there was a significant reduction in medio-lateral (F(1, 14) =14, p=.002, partial η^2^=.50) and anterior-posterior (F(1, 14) =8.8, p=.01, partial η^2^=.38) center of pressure velocity (Table 2, S3 Fig.). We did not find any differences between groups (i.e., main effect of training type) for the single-leg stance task. There were no significant differences between test-trials or training type in center of pressure kinematics in the tandem stance task (Table 3).

**Table 2.** Single-Leg Stance Center of Pressure Measures. Single-leg stance medio-lateral (ML) and anterior posterior (AP) mean center of pressure (COP) sway excursion, standard deviation (STD) of COP sway excursion, COP velocity, and COP sway area before (Pre) and after (Post) the balance beam training for 8 participants. Values are in cm, cm/s, or cm^2^. Significant p-values are denoted in bold with an asterisk.

| COP Measures (cm) | Single-Leg Stance |  |  |  |  |  |
| --- | --- | --- | --- | --- | --- | --- |
| | Unperturbed Vision | | Visual Occlusions | | p-value (partial $\eta^2$) | |
|  | Pre | Post | Pre | Post | Test-Trial | Training |
| ML sway<br>± | 0.13<br>0.05 | 0.20<br>0.12 | 0.45<br>0.43 | 0.33<br>0.48 | 0.73<br>(0.009) | 0.15<br>(0.14) |
| AP sway<br>± | 0.47<br>0.09 | 0.48<br>0.10 | 0.67<br>0.34 | 0.57<br>0.30 | 0.11<br>(0.17) | 0.23<br>(0.10) |
| STD ML sway<br>± | 0.21<br>0.08 | 0.18<br>0.05 | 0.15<br>0.04 | 0.05<br>4.54 | 0.10<br>(0.18) | 0.22<br>(0.11) |
| STD AP sway<br>± | 0.19<br>0.02 | 0.18<br>0.07 | 0.21<br>0.06 | 0.19<br>0.08 | 0.22<br>(0.11) | 0.63<br>(0.02) |
| ML velocity<br>± | 1.95<br>0.78 | 1.57<br>0.55 | 1.47<br>0.47 | 1.29<br>0.47 | **0.002\***<br>(0.50) | 0.20<br>(0.12) |
| AP velocity<br>± | 1.44<br>0.38 | 1.30<br>0.40 | 1.57<br>0.54 | 1.40<br>0.58 | **0.01\***<br>(0.38) | 0.64<br>(0.02) |
| Area<br>± | 0.74<br>0.48 | 0.63<br>0.45 | 0.57<br>0.32 | 0.61<br>0.45 | 0.37<br>(0.06) | 0.64<br>(0.02) |

**Table 3.** Tandem Stance Center of Pressure Measures. Tandem stance medio-lateral (ML) and anterior posterior (AP) mean center of pressure (COP) sway excursion, standard deviation (STD) of COP sway excursion, COP velocity, and COP sway area before (Pre) and after (Post) the balance beam training for 8 participants. Values are in cm, cm/s, or cm^2^. No significant differences were observed.

| COP Measures (cm) | Tandem Stance |  |  |  |  |  |
| --- | --- | --- | --- | --- | --- | --- |
| | Unperturbed Vision | | Visual Occlusions | | p-value (partial $\eta^2$) | |
|  | Pre | Post | Pre | Post | Test-Trial | Training |
| ML sway | 0.16 | 0.15 | 0.29 | 0.26 | 0.58 | 0.28 |
| ± | 0.11 | 0.08 | 0.32 | 0.29 | (0.02) | (0.81) |
| AP sway | 0.52 | 0.48 | 0.62 | 0.63 | 0.57 | 0.24 |
| ± | 0.15 | 0.10 | 0.29 | 0.27 | (0.02) | (0.10) |
| STD ML sway | 0.15 | 0.15 | 0.11 | 0.12 | 0.94 | 0.08 |
| ± | 0.05 | 0.04 | 0.04 | 0.03 | (0.00) | (0.20) |
| STD AP sway | 0.18 | 0.16 | 0.21 | 0.19 | 0.84 | 0.59 |
| ± | 0.12 | 0.04 | 0.04 | 0.06 | (0.003) | (0.02) |
| ML velocity | 1.34 | 1.23 | 0.90 | 0.88 | 0.43 | 0.06 |
| ± | 0.45 | 0.54 | 0.35 | 0.29 | (0.05) | (0.23) |
| AP velocity | 0.57 | 0.44 | 1.23 | 1.20 | 0.67 | 0.60 |
| ± | 0.54 | 0.20 | 0.43 | 0.50 | (0.01) | (0.02) |
| Area | 0.57 | 0.44 | 0.33 | 0.38 | 0.69 | 0.26 |
| ± | 0.54 | 0.20 | 0.19 | 0.25 | (0.01) | (0.09) |

## Discussion

Both groups showed a significant reduction in step-offs after the beam walking training. The visual occlusions training led to a higher reduction of step-offs (75%) than the unperturbed vision training (23%). Both within subjects (test trial) and between subjects (training type) effects sizes were large (η^2^=~0.7 and η^2^=0.5, respectively). Thirty minutes of treadmill-mounted beam walking practice had a large effect on motor performance, and the intermittent visual occlusions had a large effect on boosting the improvement.

The neuromechanism responsible for visual occlusion training improvement could come from multiple pathways. Previous studies have suggested that perturbing one sensory modality can enhance sensitivity and processing of other available sensory information (Assländer & Peterka, 2014, 2016; Peterka, 2002). Decreasing surface support during balance can downregulate somatosensory processing and increase reliance on visual input (Lubetzky-Vilnai et al., 2015; Song et al., 2018). However, vision is inherently coupled with processing delays and not optimal to inform postural adjustments during a challenging balance task (Stepan, 2009). Expert athletes and gymnasts tend to rely less on vision for postural control compared to novices in a wide range of sports and activities (Croix et al., 2010; Nagy et al., 2004; Paillard & Noé, 2006). Robertson and colleagues (1994), observed that experts walking on a balance beam relied more on proprioceptive and vestibular information compared to visual feedback (Robertson et al., 1994). The above studies suggest that perturbing vision by inducing re-occurring intermittent visual occlusions could have increased reliance on other sensory information (vestibular and/or proprioceptive). Alternatively, the visual occlusion could have induced neural cross-modal synchronization, a phenomenon that occurs when different sensory electrocortical oscillations synchronize due to an external stimulus (Bauer et al., 2020; Thorne et al., 2011; Voloh & Womelsdorf, 2016). Cross-modal synchronization within proprioceptive, vestibular, and visual areas, could have enhanced sensory processing and multisensory integration related to the balance task (Lakatos et al., 2007, 2008; Schroeder & Lakatos, 2009). Future studies could examine these possibilities using electroencephalography.

There were not many sacral movement parameters that changed with practice. The maximum sacral mediolateral velocity decreased with training in the visual occlusions group and the maximum sacral vertical velocity showed a change with training for both groups (Table 1, S1 Fig.). The decrease in sacral mediolateral velocity could reflect better balance control (Chiovetto et al., 2018; Huber et al., 2020). The increase in maximum sacral vertical velocity could be an indicator of more confident stepping patterns, returning to the vertical oscillations of the CoM that are characteristic to normal human walking (Kuo & Donelan, 2010; Inman & Eberhart, 1953). We did not find changes in sacral marker standard deviation in any direction. There are no consistent findings in literature regarding sacral marker variability of increased balance control during beam walking. Domingo and colleagues (2009, 2020) showed that increased sacral variability in the mediolateral direction during beam training correlated with better performance on the beam after training (Domingo & Ferris, 2009, 2010). However, no differences in mediolateral sacrum variability were observed before and after the training. Subjects did walk with a slightly shorter effective limb length after training, about 1 cm less distance between the sacral marker and ankle marker (Table 1, S2 Fig.). The lack of change in other sacral parameters during beam walking with practice suggests there was not a clear, consistent strategy in sacral movement dynamics that the subjects adopted to reduce beam step-offs.

We found limited transfer of balance beam performance gains to treadmill walking margin of stability (Fig. 3) and single-leg stance center of pressure velocity (Table 2). These measures had lower effect sizes (partial η^2^<0.5) compared to step-offs off the beam (partial η^2^>0.5), supporting the task specificity of balance training principle (Kümmel et al., 2016). Kümmel and colleagues (2016), reviewed the transfer effects of six balance training protocols in healthy participants and found limited or no balance gain transfers on non-trained tasks. However, the metaanalysis only included two studies of walking balance as the trained task (slackline walking) that consisted of multiple sessions over several weeks (Donath et al., 2013, 2016), while our study only consisted of a 30-minute training session. More recent studies looked at transfer effects of repeated slackline training sessions on other static and dynamic balance tasks and found similar results (Donath et al., 2017; Giboin et al., 2018). The above studies didn’t indicate any specific transfer effects between feed-forward and reactive balance tasks either. Future studies need to test transfer effects of a balance beam training protocol consisting of multiple training sessions on different balance tasks, including static, dynamic, and reactive balance tasks to investigate if beam walking could be used to improve balance in multiple motor tasks.

There were several limitations to the study that should be considered for future work. The sample size was relatively small. The number of subjects had been chosen based on the expected effect size of the improvements in step-offs. Indeed, we had a large effect size for the training type (~0.5) and for the pre- to post-training (~0.7) in regard to the primary performance metric (beam step-offs). The effect sizes for the sacral kinematic measures that were significantly different after beam practice were equal to or less than 0.35. Future studies that want to determine if there were small changes in sacral or CoM movement dynamics that we did not detect would need to recruit a larger sample to account for the smaller effect sizes. Another limitation of our study was that we only looked for differences in sacral movement patterns and transfer to other balance tasks after one 30-minute training session. It is possible that many sessions of training over days or weeks would have a larger effect on sacral movement dynamics and transfer to other balance tasks. We chose the time of training for this study based on the expected large effect size from 30 minutes of practice. Symeonidou and Ferris (2022) showed that 30 minutes of training would have a large effect on beam walking balance performance and that the visual perturbation training would increase the effect considerably (Symeonidou & Ferris, 2022). Future studies should extend the training time across days and weeks, to see if there is greater transfer to a wider range of balance tasks and to balance impaired patient populations. Determining if individuals of older age or with balance disorders perform similarly as the young, healthy subjects in this study would be valuable for future clinical interventions.

## Conclusion

Our results support the benefits of treadmill-mounted beam walking and intermittent visual occlusions as means to improve dynamic balance in healthy, young adults. There were few indicators of changes in sacral movement dynamics after training that may have been responsible for the large improvements in beam walking performance after half an hour of practice. Balance performance during single-leg stance and treadmill walking showed some limited transfer effects after beam walking practice, but the overall transfer was not large. Future studies examining the effects of intermittent visual occlusions using longer training interventions with multiple motor tasks on balance-challenged individuals are warranted.

## Acknowledgments

We would like to thank Evdokia Ptitsyna and Lourdes Bernandez for their assistance in data collection as well as Crosman Cruz for his assistance in data analysis.

## Author Contributions

E-RS and DF co-designed this study. E-RS acquired and analyzed the data and drafted the manuscript. NS acquired data and contributed analysis tools. RR acquired data and contributed analysis tools. DF contributed to data interpretation and manuscript drafting. All authors have read and approved the final manuscript. No one who qualifies for authorship has been omitted. The manuscript has not been published nor accepted for publication elsewhere.

## Declaration of Interests

The authors declare no competing interests. This research was supported by the U.S. National Institutes of Health (R01NS104772).

## Notes

### Competing Interest Statement

The authors have declared no competing interest.

