## Supplemental 1 Figure, Supplemental 2 Figure, Supplemental 3 Figure for "Practice walking on a treadmill-mounted balance beam modifies beam walking sacral movement and alters performance in other balance tasks"

### Supporting Information

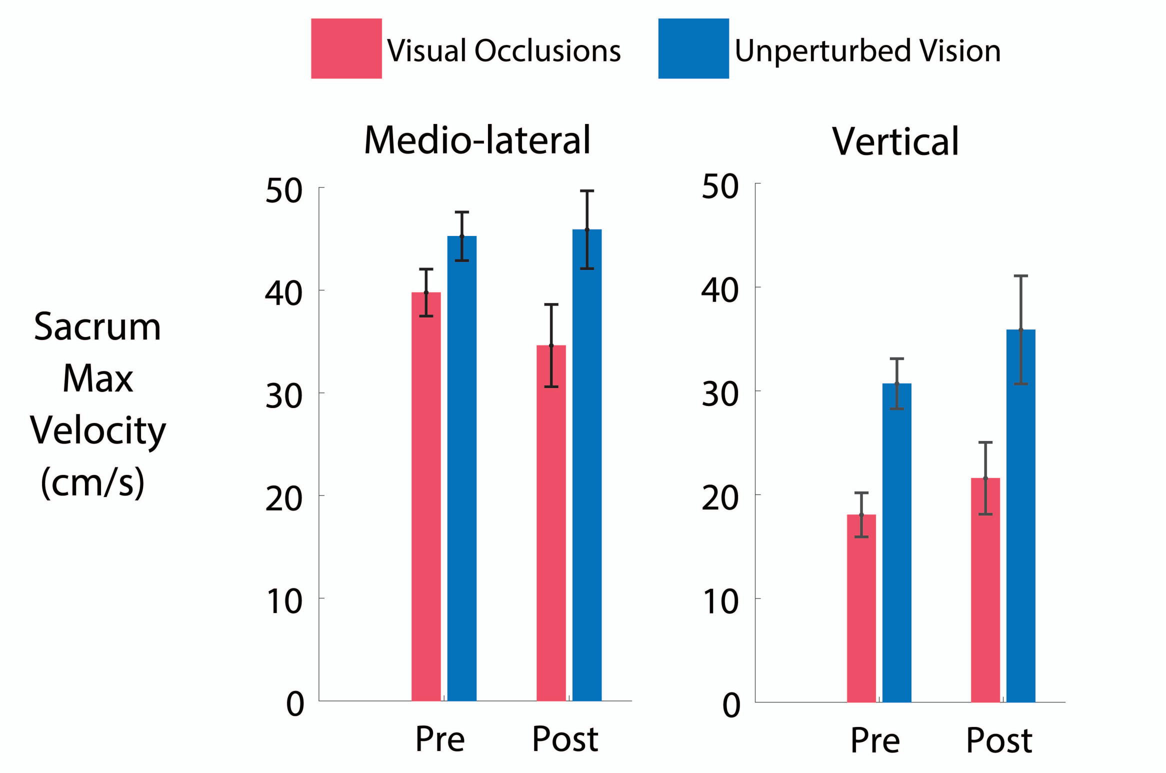


**S1 Fig. Sacral Marker Maximum Velocity.** Medio-lateral and vertical sacral marker maximum velocity for the visual occlusions (red) and unperturbed vision group (blue) before (Pre) and after (Post) the training. In the medio-lateral direction there was a significant main effect of training type ($F(1,16)=5.1$, $p=.04$, partial $\eta^2=.24$). The visual occlusions group also showed a decline in maximum sacrum velocity between pre-test and post-test. In the vertical direction we saw a main effect of training type ($F(1,16)=8.7$, $p=.009$, partial $\eta^2=.35$) and a main effect of test-trial ($F(1,16)=4.9$, $p=.04$, partial $\eta^2=.24$). Error bars represent standard error of the mean.

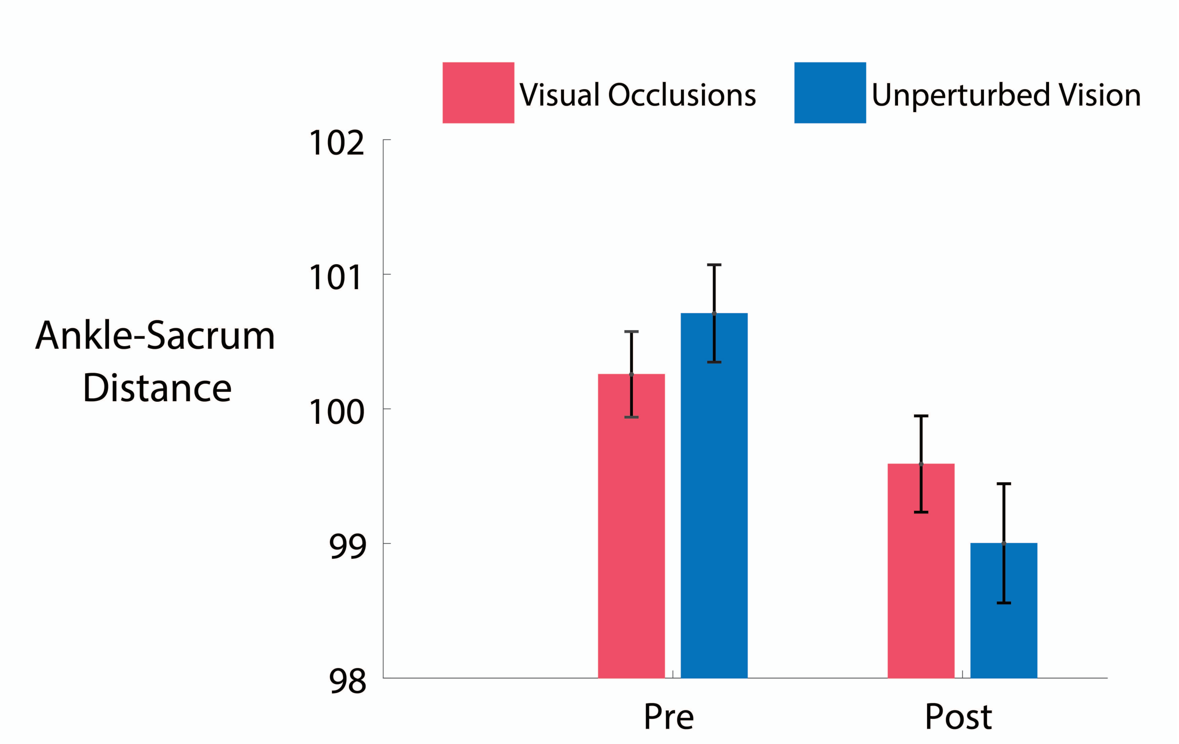


**S2 Fig. Normalized Ankle-Sacrum Distance.** Normalized ankle-sacrum distance for the visual occlusions (red) and unperturbed vision group (blue) before (Pre) and after (Post) the training. Both groups showed a significant reduction of ankle-sacrum distance between pre-test and post-test ($F(1,16)=15.8$, $p=.001$, partial $\eta^2=.5$), meaning both adapted a more crouched position after training. The visual occlusions group showed ~0.7 cm reduction, while the unperturbed vision group showed a larger deduction of ~1.7 cm. Error bars represent standard error of the mean.

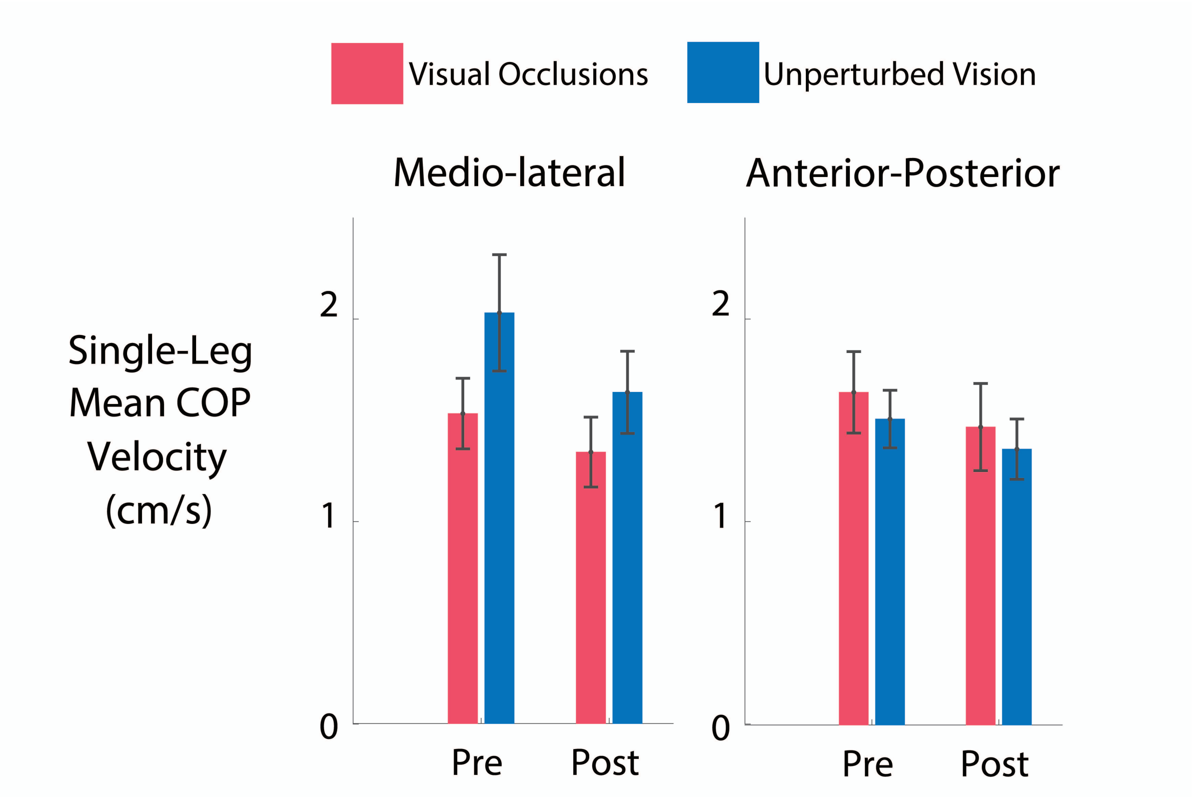


**S3 Fig. Single-Leg Stance Center of Pressure Velocity.** Mean mediolateral and anterior-posterior center of pressure (COP) velocity during single-leg stance. Both groups showed a decrease in COP velocity in the medio-lateral and anterior-posterior direction. There was a significant main effect of test-trial in the medio-lateral ($F(1, 14) = 14$, $p = .002$, partial $\eta^2 = .50$) and anterior-posterior direction ($F(1, 14) = 8.8$, $p = .01$, partial $\eta^2 = .38$). Error bars represent standard error of the mean.
